# Intrauterine inflammation leads to some sex and age-specific behavior and molecular differences in mice

**DOI:** 10.1101/2022.04.24.486005

**Authors:** Ana G. Cristancho, Natalia Tulina, Lauren Anton, Guillermo Barila, Michal A. Elovitz

**Author notes:** corresponding author Corresponding Author Contact Information: Ana G. Cristancho, MD, PhD, Abramson Research Center, Rm. 516g, Children’s Hospital of Philadelphia, 3615 Civic Center Blvd., Philadelphia, PA. 19104, U.S.A.

## Abstract

**Background:** Prenatal inflammation is associated with long-term adverse neurobehavioral outcomes in exposed children. Sex-specific differences in behavior have been observed in anxiety and learning; however, whether these differences manifest differently by age is unknown. This study assesses possible behavioral changes from *in utero* inflammation as a function of age in neonatal, juvenile, and adult animals. We also tested if the observed behavioral differences correlated to neonatal sex-specific neurogenesis gene expression changes in the hippocampus to suggest a mechanism for observed behavioral differences.

**Methods:** CD-1 timed pregnant dams were injected in utero with lipopolysaccharide (LPS, 50μg/animal) or saline at embryonic day 15 (E15). Neonatal behavioral testing was performed on postnatal day (P) 5 on male and female pups born at term using the Ultrasonic Vocalization test (USV). Juvenile and mature animals of each sex were tested in Open Field (OF) and Barnes Maze (BM) on P28 and P67 (OF), and 32-36 and 70-75 (BM). A commercially available array designed to assess the expression of genes involved in mammalian neurogenesis was utilized for profiling gene expression in the hippocampal tissue isolated from LPS and saline-exposed P7 pups.

**Results:** There were no differences in stress responses measured by neonatal USV between LPS- and saline-exposed groups of either sex. In contrast, exposure to prenatal inflammation caused a male-specific increase in anxiety in mature but not juvenile animals. Juvenile LPS-exposed females had decreased movement in OF that was not present in adult animals. In addition, we observed improved memory retrieval in response to *in utero* LPS in the juvenile animals of both sexes. However, there was an impairment of long-term memory in adult LPS-exposed females. Finally, gene expression analyses revealed that LPS induced sex-specific changes in genes involved in hippocampal neurogenesis.

**Conclusions:** Intrauterine exposure to inflammation has age and sex-specific effects on anxiety and learning. These differences are not apparent in the neonatal period but begin to be evident in juvenile animals and evolve in adult animals. These sex-specific differences in learning may be correlated to sex-specific disruption of the expression of genes associated with neurogenesis in the hippocampus.

## BACKGROUND

Clinical data strongly support the link between maternal infections during pregnancy and a wide range of adverse neurobehavioral outcomes in exposed children, such as learning delay, cerebral palsy, depression, anxiety, schizophrenia, epilepsy, attention-deficit disorder, and autism [1-4]. These neurological abnormalities from intrauterine inflammation are thought to result from cellular and molecular alterations that lead to grey and white matter injury in the developing brain [5-8]. Experimental animal models have suggested that activation of the innate immune response causes long-lasting changes in brain function and behavior [9, 10]. In models of maternal systemic inflammation, previous studies have shown that the activation of the Toll-like receptor (TLR) signaling pathway during pregnancy leads to alterations in certain behavioral outcomes [6-14]. However, systemic maternal inflammation is only one mechanism by which prenatal inflammation can occur and it does not recapitulate the local inflammation that is frequently present in preterm birth [15, 16].

Here we use a mouse model of localized intrauterine inflammation to better recapitulate a common clinical scenario by which a fetus is exposed to inflammation [16]. This model has been shown to produce molecular and cellular hallmarks of fetal and postnatal brain injury [16-20]. With this model, we can assess possible behavioral alterations resulting from prenatal infection and inflammation which were induced by *in utero* exposure to a sterile inflammatory stimulus (lipopolysaccharide, LPS). Using this model, we previously reported that exposure to intrauterine inflammation did not result in sex-specific differences in anxiety in 12-14 week old mice [21]. However, recent evidence demonstrates that sex and age both independently and synergistically alter anxiety-related behaviors and learning, with significant differences sometimes observed in juvenile mice that are absent in adult animals [22-25]. There are some sex-specific differences in the early neurodevelopmental milestone of negative geotaxis, where males are slower to turn around when placed on an incline in the first 10 days of life [26].

While these studies demonstrate sex-specific behavioral development patterns, there have been no studies investigating the effects of intrauterine inflammation with sex throughout the age span on anxiety and learning. Here we compare the development of stress-induced anxiety response and animals’ performance in spatial learning and memory tasks in the offspring of dams exposed to in utero inflammation or saline during pregnancy. Since anxiety related behaviors and learning and memory are known to be influenced by changes in hippocampal neurogenesis [27-31], we also determined if the expression levels of genes associated with adult neurogenesis were altered in response to LPS exposure in male and female pups. A better understanding of how prenatal inflammation affects behavior in exposed offspring in a sex-specific manner and over a lifetime is important for developing new therapeutic approaches to improve inflammation-induced adverse neurobehavioral outcomes.

## METHODS

### Mouse model of intrauterine inflammation

An established mouse model of intrauterine inflammation [16] was used to study the effect of prenatal immune activation on specific behavioral outcomes in exposed offspring. Briefly, a mini-laparotomy was performed on CD-1 timed pregnant mice on E15. A single injection of *E. coli* LPS (055:B5, Sigma Aldrich, St. Louis, MO; 50μg/100μl per animal) or saline (100μl per animal) was applied to the right intrauterine horn between first and second gestational sacs. For Open Field (OF) and Barnes Maze (BM) experiments the pups delivered at term were culled to three pups per sex on P2-P3 and weaned on P21. Animals were maintained in a 12:12 hr light:dark cycle and were allowed to feed *ad libitum*.

All animal care and treatment procedures were approved by the Institutional Animal Care Committee following the guidelines from the National Institutes of Health.

### Behavioral Testing

Animal behavior was assessed in pups born at term using the following behavioral tests: Ultrasonic Vocalization (USV), Open Field (OF) and Barnes Maze (BM). For USV testing 11 male and female pups from different litters were tested per group (LPS or saline) on P5. The OF test was done using preadolescent (P28) and mature (P67) pups of both sexes from LPS- and saline-exposed groups. One male and one female pup were tested per group per litter: LPS-treated males: n=13; LPS-treated females: n=13; saline-treated males: n=12; saline-treated females: n=13. The preadolescent and mature pups tested in OF were subsequently subjected to BM testing on P32-36 and P70-74, respectively.

The USV test was performed on the offspring of saline and LPS injected dams at P5 according to a previously published protocol [32]. Briefly, USVs were recorded using an UltraSoundGate Condenser Microphone CM 16 (Avisoft Bioacoustics, Berlin, Germany) which was placed 12 cm above the testing surface within sound attenuating cabinet. The recordings were made with Avisoft Recorder software (version 4.2.14; Avisoft Bioacoustics, Glienicke, Germany) using a sampling rate of 375,000 Hz in a 16-bit format. For signal analysis, fast Fourier transformation was performed with Avisoft SASLab Pro (version 4.40; Avisoft Bioacoustics, Glienicke, Germany) to generate spectrograms. Spectrograms were visually scanned to remove artifacts and the number of ultrasonic vocalizations (calls), average call duration and the frequency of emitted sound was determined during a 5 minute recording session.

To determine the effect of prenatal inflammation on exploration and anxiety responses, the OF testing was performed on preadolescent and mature pups of both sexes. At the beginning of each 10 min. trial mice were placed individually at the center of a round-shaped arena (San Diego Instruments, San Diego, CA). A scaffold of infrared emitters and photodetectors was placed around the arena to detect beam breaks as the mouse moved. XY-axis detectors collected peripheral and center beam breaks as their peripheral and central activities, respectively, and an elevated Z-axis detector collected vertical beam breaks as rearings. Rearings as well as central, peripheral, horizontal (the sum of central and peripheral activities) and total (the sum of rearings and horizontal activity) activities were recorded using the PAS-Open Field system (San Diego Instruments, San Diego, CA). While most of these measurements were presented as the average number of beam breaks per a 10 min. trial, central activity was expressed as a percentile from horizontal activity: (center activity/ horizontal activity) x 100%.

To determine possible differences in spatial learning and memory between LPS- and saline-exposed groups, we conducted BM testing on male and female pups from both groups. BM was made out of a large, flat disc with twenty holes at the perimeter (San Diego Instruments, San Diego, CA). All holes, except one, escape, hole, were blocked during acquisition trials. Landmark cues with unique geometric shapes were placed around the maze which was illuminated to ∼1000 lux from an overhead halogen lamp. The escape hole was demarked by a specific visible cue. All trials were recorded using Microsoft Lifecams (Microsoft, Redmond, VA) for documentation and offline analysis. The procedure consisted of four acquisition sessions performed on four consecutive days, specifically, P32-35 and P70-73 for preadolescent and mature animals, respectively, followed by a probe trial after acquisition. Two acquisition trials per day were done with a 30min. interval. If the escape compartment was entered, the trial ended and the latency to escape was recorded. If a mouse did not enter the escape compartment in 150 seconds, it was gently guided into the hole.

Twenty-four hours after the last acquisition trial, on P36 and P74, respectively, a 2-minute probe trial was performed with all holes occluded, including the escape compartment. Image analysis software (ANYmaze, Stoelting Co., Wood Dale, IL) was used to divide the maze into four equal quadrants virtually, and the time spent in each quadrant was recorded. Mice that learned to associate the cued-quadrant with the escape compartment location spent more time in the quadrant where the cue was located.

### Hippocampal dissections

Hippocampal dissections were performed on P7 brains isolated from male and female offspring of LPS and saline-treated dams as published [33].

### Gene expression analysis using Mouse Neurogenesis Arrays

Prior to analyzing possible changes in gene expression associated with mouse neurogenesis, the sex of each pup was determined by quantifying male- and female-specific transcripts, SRY and XIST, respectively, in RNA samples isolated from pups’ tails. Tissue homogenization was achieved using TissueLyser II (QIAgen Sciences, Germantown, MD) and a standard protocol for RNA extraction with QIAzol Lysis reagent (QIAgen Sciences, Germantown, MD) and 1-bromo-3-chloropropane reagent (BCP, Thermo Fisher Scientific, Philadelphia, PA) was applied. After assigning information about sex for each sample all samples were subdivided into the following four groups: LPS-treated female samples (n=13), LPS-treated male samples (n=10), saline-treated female samples (n=5) and saline-treated male samples (n=5).

Subsequently, total RNA was isolated from the hippocampi of male and female pups exposed to prenatal LPS or saline using the RNeasy Microarray Tissue mini kit (QIAgen Sciences, Germantown, MD). The integrity of RNA samples was confirmed using 2100 Bioanalyzer (Agilent Genomics, Santa Clara, CA) and cDNAs were synthesized using the RT^2^ First Strand Kit (Qiagen Sciences, Germantown, MD). Gene expression was compared between the hippocampal samples from LPS- and saline-exposed pups using Mouse Neurogenesis Arrays (RT^2^ Profiler PCR Array, Qiagen Sciences, Germantown, MD). Expression levels of 84 genes with well-documented roles in murine neurogenesis were examined. qPCR amplification was done using RT^2^ SYBR Green ROX qPCR Mastermix (Qiagen Sciences, Germantown, MD). All RT-PCR reactions were repeated four times and changes in gene expression were analyzed using the ΔΔC_T_ method.

Differentially expressed genes for each sex were entered into GeneMania to determine functional networks regulated by intrauterine inflammation in the hippocampus [34] (https://genemania.org, blue version). The top 5 pathways were noted for each sex.

### Statistical Analysis

Statistics were performed as described previously using R-studio [35]. Briefly, we used mixed models with generalized estimating equations in the geepack package in R with cohort as the random effect [36]. Linear mixed-effects models in the lme4 package in R was used for BM since repeated measures between trials were required [37]. Statistical significance is shown for LPS-exposure, sex, and interaction between LPS-exposure and sex since intrauterine inflammation has been shown to have sex-dichotomous effects [21]. Statistical significance was set at * *p* < 0.05, ** *p* < 0.01, and *** *p* < 0.001. We corrected *p*-values from mixed model analyses with a Benjamini-Hochberg correction.

Gene expression data generated using Mouse Neurogenesis Arrays were analyzed on the RT^2^ Profiler PCR Array Data Analysis Webportal (QIAgen Sciences, Germantown, MD) using the ΔΔC_T_ method. Target genes were normalized to three housekeeping genes: β-Glucuronidase (*Gusb*), β-2 microglobulin (*B2m*), and Heat shock protein 90 α (cytosolic), class B member 1 (*Hsp90ab1*). The replicate 2^(-ΔCT)^ values were compared for each target gene in LPS and saline treated groups using Student’s T-test (two-tail distribution with equal variances between the two groups of samples). We corrected *p*-values from mixed model analyses with a Benjamini-Hochberg correction. Male and female datasets were analyzed separately. The corrected *p*-values that are less than 0.10 were considered significant. The fold change in gene expression levels between LPS and saline treated groups were calculated as 2^(-ΔΔCT)^.

## RESULTS

### Intrauterine inflammation does not alter ultrasonic vocalizations in neonatal pups

USV testing in the neonatal period can study the pups’ response to the stress of separation from their mothers and as an assessment of early deficits in communication [38]. Anxiety-related phenotypes are associated with increased vocalizations, whereas autism-related phenotypes are associated with decreased vocalizations [38]. Therefore, we performed USV testing on saline- and LPS-exposed male and female pups on P5 isolated from their mothers as an early test for neurodevelopmental deficits. Vocal responses resulting from the isolation from mother, specifically, the total number of calls per 5-minute interval **(Fig. 2A)** and the frequency of emitted sound **(Fig. 2B)**, were compared between the groups. Our data show no significant changes in vocal responses between saline and LPS groups in male or female animals. Thus, LPS exposure does not affect social communication skills, at least at this early stage of postnatal development.

**Figure 1:**
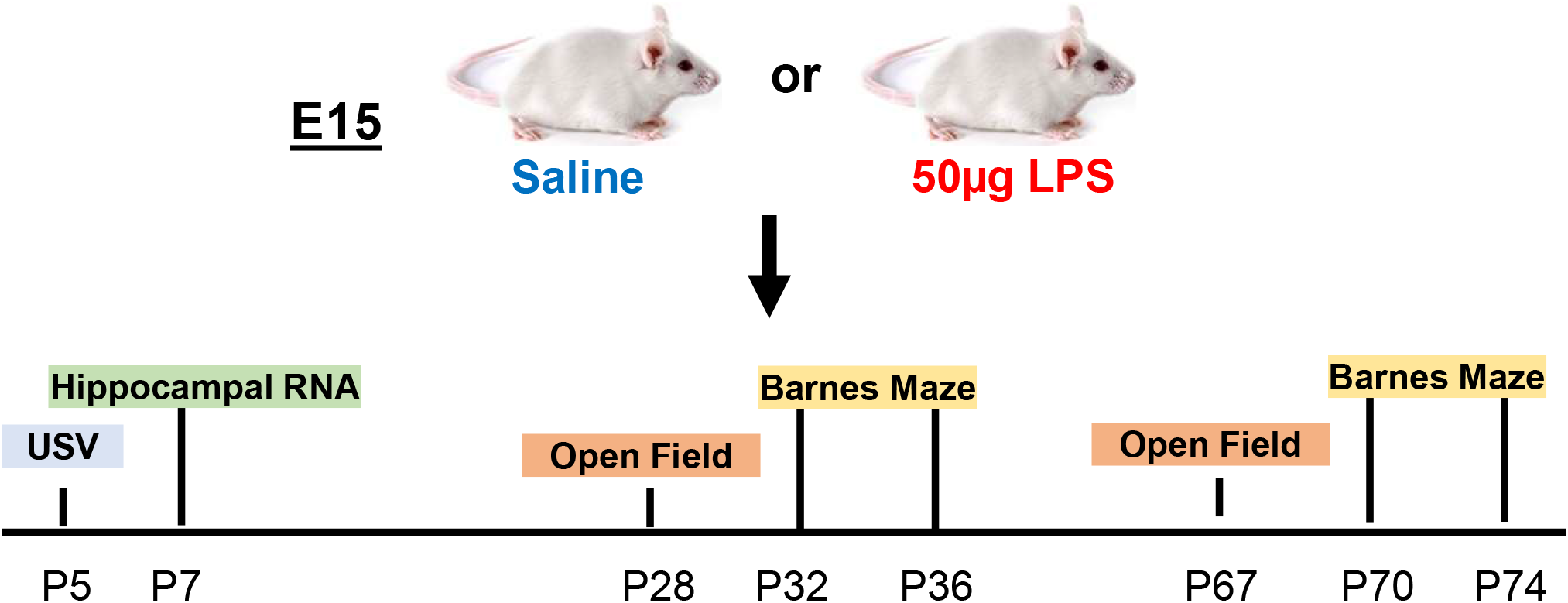
Schematic and timeline of experiments.

**Figure 2:**
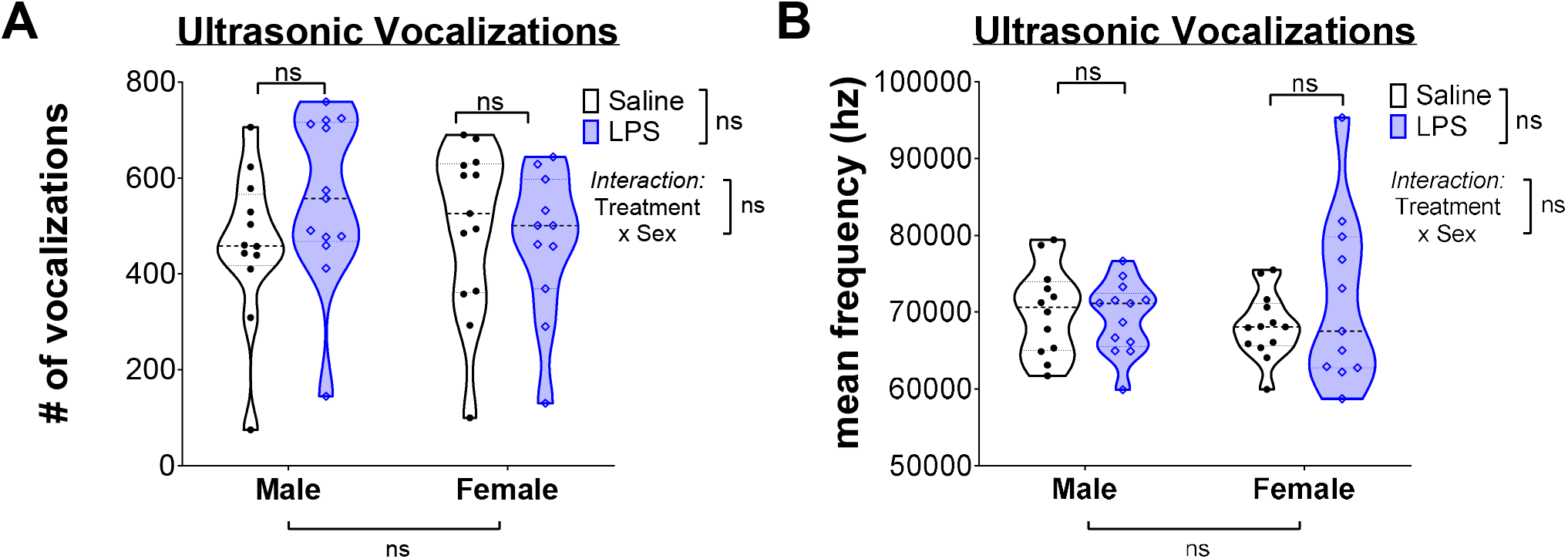
Intrauterine inflammation is not correlated to changes in USV. Violin plots quantifying the number (A) and frequency (B) of USV. Individual points represent single animal.

### Intrauterine inflammation leads to increased anxiety-related behavior in adult, but not juvenile, males

We conducted OF testing to determine if intrauterine inflammation altered exploratory behaviors in a new environment in juvenile, preadolescent (P28) and mature (P67) animals. Decreased exploration of the center of an open field is associated with increased anxiety-related deficits [39-41]. LPS-exposure did not increase time spent in the center of an open field for either sex in juvenile animals **(Fig. 3A)**. However, female juvenile mice exposed to LPS had decreased horizontal activity **(Fig. 3B)**, reflective of a possible subtle motor deficit. By contrast, LPS-exposure significantly reduced the time male adult mice spent in the open field center, consistent with increased anxiety **(Fig. 3C)**. Neither sex had deficits in motor activity from LPS **(Fig. 3D)**, consistent with the resolution of the motor deficit observed in juvenile female mice from LPS.

**Figure 3:**
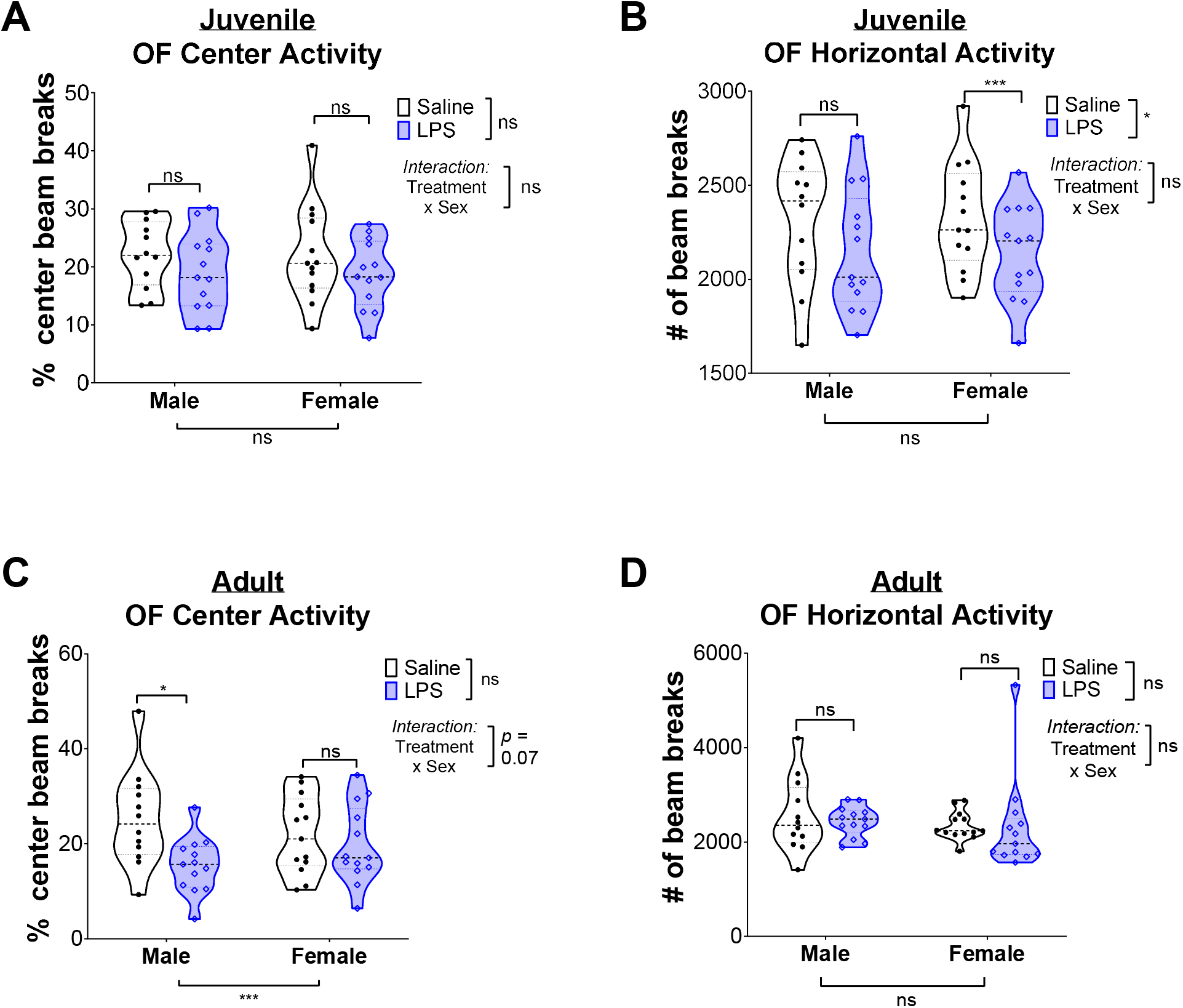
Adult males have increased anxiety-like behaviors after exposure to intrauterine inflammation. Violin plots quantifying the (A) percent of center beam breaks in open field and the (B) extent of total movement in juvenile animals in OF. In (C-D) these same parameters are quantified in adult animals. Individual points represent single animal.

### Intrauterine exposure to inflammation is not associated with improved memory in juvenile mice, but impaired memory in adult female mice

We next sought to test whether exposure to intrauterine inflammation predisposed animals to spatial learning and memory deficits. We used the BM test in juvenile (P32-36) and adult (P70-P74) animals. Male and female pups were tested as separate groups. Spatial learning is tested by using visual cues to direct the animals towards an “escape hole” in the apparatus. As expected, over the course of the acquisition trials, the male and female animals of both age groups were able to escape the apparatus more quickly, reflecting typical spatial learning **(Fig. 4A-D)**. There was no difference in the rate of learning between saline-exposed and LPS-exposed animals.

**Figure 4:**
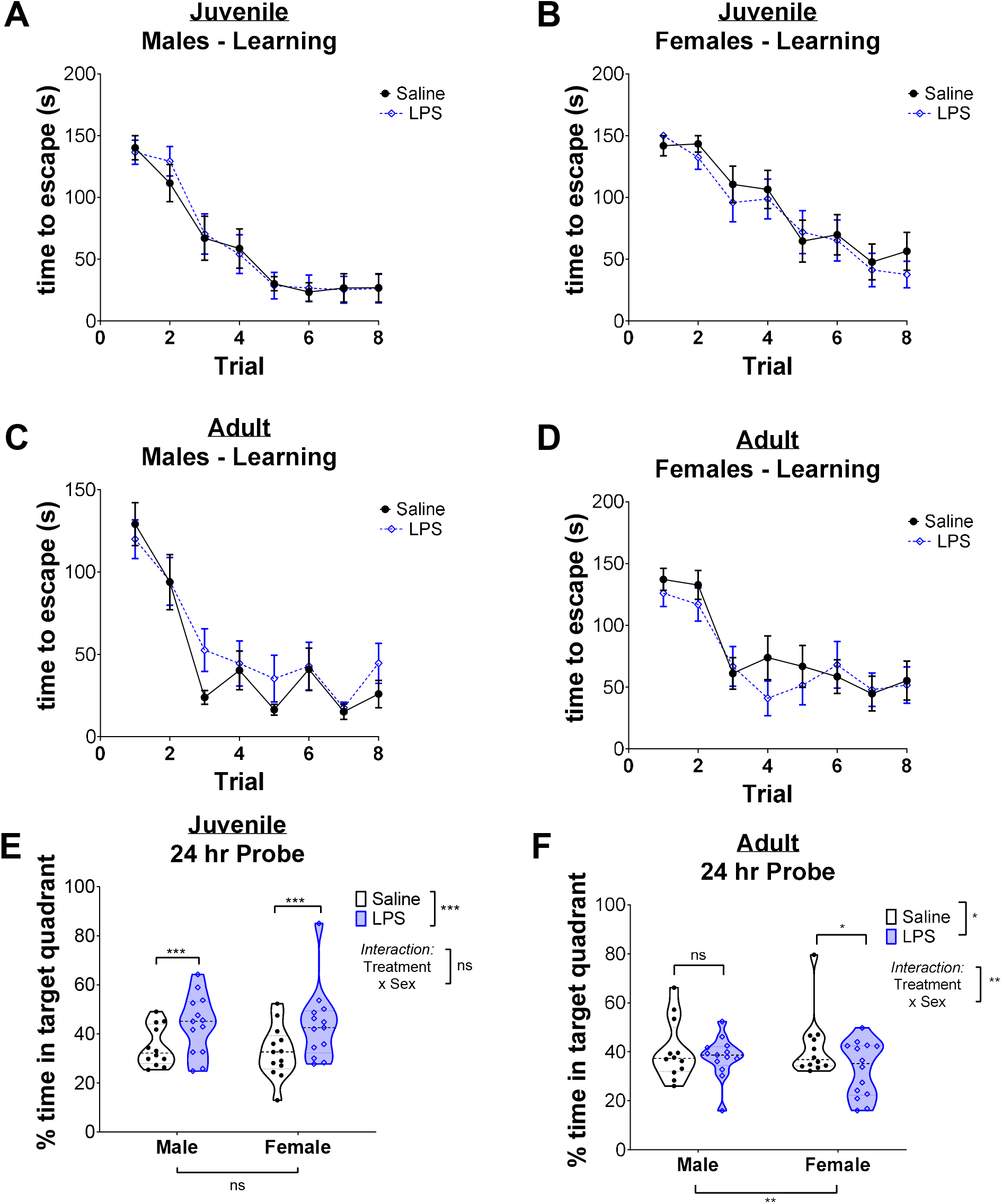
Intrauterine inflammation leads to accentuation of memory in juvenile animals and a disruption of memory in adult female animals. (A-D) Time to escape in juvenile and adult animals. (E-F) Percent time spent in target quadrant for long-term memory recall studies in juvenile and adult mice. Individual points represent single animal.

However, substantial differences were found in the twenty-four hour probe trial that may suggest differences in memory. Surprisingly, male and female juvenile mice, exposed to intrauterine inflammation, spent more time in the target quadrant than saline-exposed animals **(Fig. 4E)**, suggesting an improvement in memory. By contrast, in adult mice, there were no differences in male mice by exposure, but female mice exposed to intrauterine inflammation had a significant decrease in time in the target quadrant **(Fig. 4F)**, suggesting a sex-dichotomous reduction in long-term memory.

### Intrauterine exposure to inflammation leads to sex-dichotomous differences in gene expression related to neurogenesis

Expression levels of 84 genes known to play important roles in mouse neurogenesis were compared in the hippocampi of saline- and LPS-exposed pups using commercial RT-PCR arrays on P7. This analysis revealed there were hippocampal transcripts altered in response to prenatal inflammation in females only, in males only or in both sexes when using an FDR <0.10 as a cut off **(Fig. 5A)**. Most of these changes were less than 2-fold, except for Amyloid β A4 precursor protein-binding (Apbb1) and Notch gene homolog 2 (Notch2) genes in intrauterine inflammation exposed females. There were 9 genes that only changed in females (white), 2 genes that only changed in males (light grey), and 1 gene, *Pafah1b1*, that changed in both sexes with LPS (dark grey). Of note, none of the genes that changed in male or female mice were different in saline-exposed mice between males and females.

**Figure 5:**
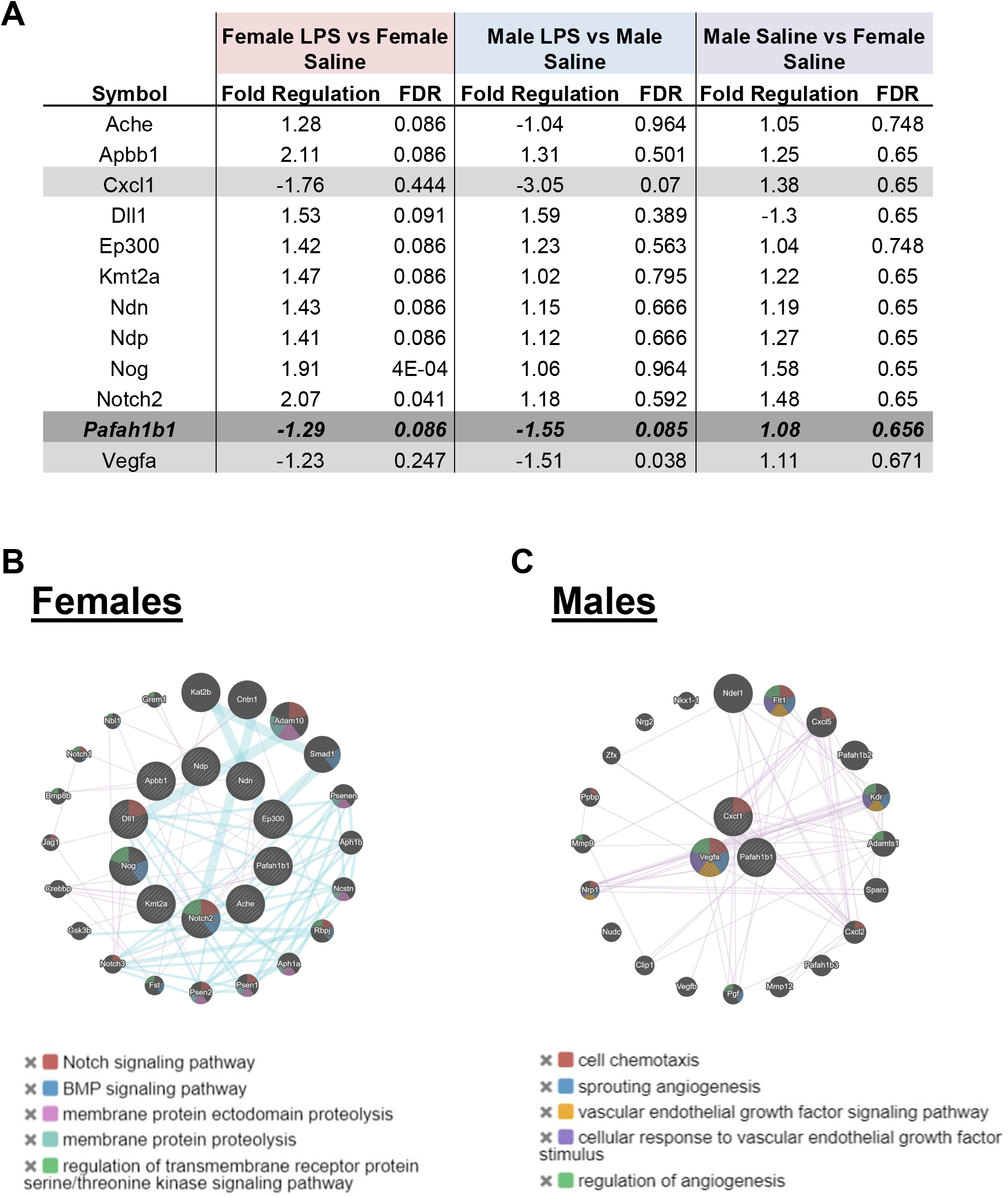
Sex-dichotomies in neonatal hippocampal gene expression from intrauterine inflammation. (A) Table of neurogenesis related genes that are dysregulated in the P7 hippocampus. Genes in white change in females only. Genes in light grey change in males in only. *Pafah1b1*, in dark grey, is the only gene that changed in both males and females. GeneMania pathway analysis in genes differentially expressed in (B) females and males (C) (genes in the inner circle). Colored shading corresponds to the top 5 pathways enriched by an FDR < 5%.

Previous research has demonstrated that intrauterine inflammation regulated neurogenesis [42]. To determine if there were sex-dichotomous neurogenesis related pathways modulated by intrauterine inflammation, we used GeneMania to build weighted interaction networks based on differentially expressed genes. Genes regulated by intrauterine inflammation in female mice were enriched for the Notch and BMP pathway **(Fig. 5B)**. By contrast, the few genes regulated in males were related to chemotaxis and angiogenesis **(Fig. 5C)**. These findings suggest there may be sex-dichotomous pathways related by intrauterine inflammation in the hippocampus that may be accounting for sex differences in behavior.

## DISCUSSION

Exposure to intrauterine inflammation is a well-known risk factor for both preterm birth and a spectrum of neurological, cognitive, and sensory-motor deficits that are manifested in exposed children during childhood and later in life [1-4]. However, relatively little is known about how age and sex of the offspring of exposed mothers affect these negative neurobehavioral outcomes. In this study, we utilized an established mouse model of intrauterine inflammation to show that *in utero* exposure to a sterile inflammation, can cause distinct behavioral changes in exposed offspring that are both sex specific and manifested differently throughout the lifetime. In particular, our data demonstrate that mature, but not pre-adolescent, male pups which were exposed to *in utero* inflammation have elevated anxiety levels when transitioned to an open space. In addition, we observed a temporary improvement in the retrieval of spatial memory in pre-adolescent pups of both sexes after exposure to inflammation; however, this behavioral change was not detected in older LPS-exposed animals suggesting age-specific evolution of memory. Indeed, female adult animals exposed to LPS had worse long-term memory. Lastly, prenatal inflammation altered expression levels of multiple genes involved in hippocampal neurogenesis. Interestingly, some altered transcripts showed sex-specific changes, while others were dysregulated in both male and female samples. These changes in expression levels are present during the early postnatal period (P7) and, therefore, may underlie, at least in part, the observed behavioral alterations.

Epidemiological studies have shown that inflammation during pregnancy is associated with a higher incidence of autism spectrum disorders in exposed children [7]. The link between prenatal inflammatory exposure and the symptoms of autism was further demonstrated by using experimental animal models in which an innate immune response was induced prenatally by injecting bacterial or viral ligands systemically [43-47]. In this study, we addressed a possible effect of localized intrauterine inflammation on the development of social communication skills during early postnatal period by using the USV test. In contrast to previous reports showing altered USV responses after systemic exposure to inflammatory stimuli [44-47], we did not reveal any differences between LPS- and saline-exposed groups in either sex. However, it should be noted that there is controversy regarding the effect of inflammation on USV responses as studies have shown both increases and decreases in vocalization after exposure to an inflammatory agent [43-47]. These discrepancies could result from the use of different mouse strains or various ages since both factors are shown to affect the behavioral performance in the USV test. For example, in the BTBR strain, exposure to prenatal inflammation led to an increase in the number of calls at P8, P10 and P12, while similar changes in vocalization response in C57/B6 mice exposed to the same inflammatory stimulus were detected only at P10 [48]. As in our study social behavior was examined in a different mouse strain (CD-1) at a single time point (P5), we may have missed possible behavioral alterations in LPS-treated pups at other time points which were not studied.

Increased anxiety is known to accompany many neurological conditions, including those linked to prenatal inflammation, such as depression, autism, epilepsy, schizophrenia and others [49, 50]. We addressed whether the effect of prenatal inflammation on stress-induced exploratory behavior and anxiety levels depends on the sex of exposed offspring when examined at two different time points throughout their lifetime. Our data, obtained using the OF test, revealed that exposure to prenatal inflammation caused increased anxiety in LPS-exposed mature (P68), but not pre-adolescent (P27), male pups, while female pups from both age groups remained unaffected. An increase in anxiety levels in LPS-exposed male pups detected in our study is consistent with previously published data which were obtained using both systemic immune activation [51-59], intrauterine exposure to infectious agents or their ligands [21, 60], and prenatal hypoxia [35]. For instance, using the OF test, Dada and co-authors showed that mouse offspring that received intrauterine LPS injections at E17 had elevated anxiety levels at P60, although a sex-specific effects were not tested [60]. Other work has shown that *in utero* inflammation leads to higher anxiety in both male and female pups between P84 and P98 [21]. These findings and our OF data demonstrate that prenatal exposure to inflammation facilitates stress-induced anxiety in both exposed male and female offspring at around the time of their sexual maturation; however, this effect is manifested earlier in males (P68, this study) than in females (P84-98) [21]. Therefore, sex and age play important roles in altering stress-induced anxiety in pups exposed to prenatal inflammation. Recent publications showing sex- and/or age-specific metabolic [61], immunological [21] and gene expression (this study, see below) changes in the offspring of inflammation-exposed dams may provide clues into possible molecular and cellular mechanisms underlying differential behavioral responses in a sex-specific manner.

We also determined if spatial learning and memory, behaviors that can be readily tested in mice, were altered in LPS-exposed pups. Our data showed that a mouse’s ability to learn and acquire spatial memories, after exposure to LPS, did not change compared to a saline control by sex or over the lifespan. In contrast, memory retrieval was moderately improved in LPS-exposed pups of both sexes at the younger age (P27). This effect was transient as it did not persist at the later time point (P68). The area of the brain responsible for spatial learning and memory is known to be localized to the hippocampus [62-64]. The execution of these behavioral tasks relies on the function of hippocampal granule neurons that are produced both before birth and throughout adulthood in the hippocampal subgranular zone (SGZ) [65]. Interestingly, silencing of adult-born granule neurons using an optogenetic approach is shown to specifically impair memory retrieval, but not memory acquisition [66], which indicates that novel neuronal production is exclusively responsible for memory retention. Furthermore, adult neurogenesis can be modulated by various environmental and physiological stimuli, including the inflammatory status of the organism [67, 68]. Our group recently demonstrated that intrauterine inflammation leads to the reduction in the rate of hippocampal neurogenesis between P7 and P14 and a subsequent decrease in total neuronal density at P28 [42]. Remarkably, these alterations in neuronal production in the hippocampal SGZ appear to take place at around the same time in postnatal development when an improvement in memory retrieval was observed in this study (P27). Previous data show that the disruptions in hippocampal neurogenesis can result in an impaired ability to form and retrieve spatial memories [69, 70], which is the opposite effect compared to our BM experiments. The difference between our results and those from previous studies could be accounted for by the fact that in this earlier work specific techniques were applied to severely reduce or even completely ablate neurogenesis, such as the use of antimitotic agents and irradiation, respectively, while in our mouse model only a modest reduction in the hippocampal neurogenesis was achieved after exposure to intrauterine inflammation [42]. Therefore, it is possible that these differential changes in neurogenesis would lead to different behavioral effects in the BM test. The mechanism by which the decrease in hippocampal neurogenesis could facilitate memory retrieval may involve an increase in electrical activity of the remaining granule cells as means of compensation for the function of missing neurons. In fact, there is evidence that hippocampal granule neurons from pups exposed to *in utero* inflammation have increased excitatory synaptic strength [71], which suggests that these neurons are excited more frequently that those from inflammation-free pups. It should, however, be noted that other brain areas and neuronal circuits may also be responsible for the modulation of memory retrieval following exposure to prenatal inflammation. For instance, it has been shown that infants who survived clinical infections prior to birth show aberrantly increased cortisol levels [72, 73]. While stress-induced cortisol response is known to both enhance and reduce memory performance [74], data show that a low level of cortisol found in aging population correlates with the decline in spatial memory [75]. Therefore, it is possible that an inflammation-induced increase in cortisol level may have an opposite effect improving memory retrieval in our animal model.

In contrast to *in utero* inflammatory stimulation, systemic maternal inflammation during pregnancy is shown to disrupt, not improve, the behavioral performance of exposed offspring in spatial learning and memory tasks [76-80]. Yet, one study demonstrated an increase in memory retrieval in mouse male offspring of dams injected intraperitoneally with staphylococcal endotoxin A [81]. This suggests that different mechanisms are involved in the regulation of these behavioral outputs dependent on how infectious agents or their ligands reach and react in the placental-fetal compartment.

Disruptions in hippocampal neurogenesis can also lead to changes in other behavioral outputs, such as stress-induced anxiety response [65]. In particular, genetically-induced alterations in neuronal production in the hippocampal dentate gyrus were shown to have an inverse effect on anxiety levels [31, 82]. Therefore, the reduction in hippocampal neurogenesis in LPS-exposed pups [42] is consistent with our finding of increased anxiety after exposure to intrauterine inflammation. In addition, our gene expression data show that prenatal inflammation can cause sex-specific changes in expression levels of various regulators of murine neurogenesis at P7. These data shed some light on possible molecular mechanisms associated with hippocampal neurogenesis which may underlie differential behavioral responses to a prenatal immune stimulation in different sexes, such as heightened anxiety observed in mature male, but not female, pups following *in utero* inflammation. Our results show that inflammation-induced alterations in hippocampal gene expression occur in several molecular pathways, including signaling cascades, such as Notch, BMP, Wnt/β-catenin, MAPK and Nrg1, transcriptional and/or chromatin regulation, neurotransmission, apoptosis and App-associated pathology among others. While most of these pathways are altered in both sexes, some of their molecular components showed changes only in male or female hippocampi. For example, among several members of the Notch signaling pathway whose levels were elevated in response to *in utero* LPS treatment, *Delta-like 1* (*Dll1*) ligand and *Notch2* receptor were increased in females. The Notch signaling pathway is known to be involved in the regulation of multiple events during adult hippocampal neurogenesis [83], therefore, these differences in the downstream responses to inflammation-induced Notch pathway activation suggest differential effects on novel neuronal production in the hippocampi of male and female pups which may in turn lead to alternative behavioral outcomes, such as those observed in this study.

In conclusion, here we present sex-dichotomies with age after intrauterine inflammation in anxiety, memory, and gene expression in the hippocampus. This work suggests that the interaction of age and sex are important to consider for behavior studies of intrauterine insults. Future studies will focus on understanding which pathways may be regulating these differences and whether the expression differences persist in adulthood to continue account for behavior differences. It is possible these pathways can be targets for therapeutic interventions to improve long-term outcomes.

## REFERENCES

1. Leung MP, Thompson B, Black J, Dai S, Alsweiler JM. The effects of preterm birth on visual development. Clin Exp Optom. 2018;101:1:4–12; doi:10.1111/cxo.12578.

2. Vieira ME, Linhares MB. Developmental outcomes and quality of life in children born preterm at preschool-and school-age. J Pediatr (Rio J). 2011;87:4:281–91; doi:10.2223/JPED.2096.

3. Allen MC, Cristofalo EA, Kim C. Outcomes of preterm infants: morbidity replaces mortality. Clin Perinatol. 2011;38:3:441–54; doi:10.1016/j.clp.2011.06.011.

4. Fawke J. Neurological outcomes following preterm birth. Semin Fetal Neonatal Med. 2007;12:5:374–82; doi:10.1016/j.siny.2007.06.002.

5. Dammann O, Leviton A. Maternal intrauterine infection, cytokines, and brain damage in the preterm newborn. Pediatr Res. 1997;42:1:1–8; doi:10.1203/00006450-199707000-00001.

6. Strunk T, Inder T, Wang X, Burgner D, Mallard C, Levy O. Infection-induced inflammation and cerebral injury in preterm infants. Lancet Infect Dis. 2014;14:8:751–62; doi:10.1016/S1473-3099(14)70710-8.

7. Knuesel I, Chicha L, Britschgi M, Schobel SA, Bodmer M, Hellings JA, et al. Maternal immune activation and abnormal brain development across CNS disorders. Nat Rev Neurol. 2014;10:11:643–60; doi:10.1038/nrneurol.2014.187.

8. Wang X, Hellgren G, Lofqvist C, Li W, Hellstrom A, Hagberg H, et al. White matter damage after chronic subclinical inflammation in newborn mice. J Child Neurol. 2009;24:9:1171–8; doi:10.1177/0883073809338068.

9. Burd I, Balakrishnan B, Kannan S. Models of fetal brain injury, intrauterine inflammation, and preterm birth. Am J Reprod Immunol. 2012;67:4:287–94; doi:10.1111/j.1600-0897.2012.01110.x.

10. Wang X, Hagberg H, Zhu C, Jacobsson B, Mallard C. Effects of intrauterine inflammation on the developing mouse brain. Brain Res. 2007;1144:180–5; doi:10.1016/j.brainres.2007.01.083.

11. Lehnardt S, Lachance C, Patrizi S, Lefebvre S, Follett PL, Jensen FE, et al. The toll-like receptor TLR4 is necessary for lipopolysaccharide-induced oligodendrocyte injury in the CNS. J Neurosci. 2002;22:7:2478–86; doi:20026268.

12. Mallard C, Wang X, Hagberg H. The role of Toll-like receptors in perinatal brain injury. Clin Perinatol. 2009;36:4:763–72, v-vi; doi:10.1016/j.clp.2009.07.009.

13. Wang X, Rousset CI, Hagberg H, Mallard C. Lipopolysaccharide-induced inflammation and perinatal brain injury. Semin Fetal Neonatal Med. 2006;11:5:343–53; doi:10.1016/j.siny.2006.04.002.

14. Yuan TM, Sun Y, Zhan CY, Yu HM. Intrauterine infection/inflammation and perinatal brain damage: role of glial cells and Toll-like receptor signaling. J Neuroimmunol. 2010;229:1–2:16-25; doi:10.1016/j.jneuroim.2010.08.008.

15. Bauman MD, Van de Water J. Translational opportunities in the prenatal immune environment: Promises and limitations of the maternal immune activation model. Neurobiol Dis. 2020;141:104864; doi:10.1016/j.nbd.2020.104864.

16. Elovitz MA, Wang Z, Chien EK, Rychlik DF, Phillippe M. A new model for inflammation-induced preterm birth: the role of platelet-activating factor and Toll-like receptor-4. Am J Pathol. 2003;163:5:2103–11; doi:10.1016/S0002-9440(10)63567-5.

17. Leitner K, Al Shammary M, McLane M, Johnston MV, Elovitz MA, Burd I. IL-1 receptor blockade prevents fetal cortical brain injury but not preterm birth in a mouse model of inflammation-induced preterm birth and perinatal brain injury. Am J Reprod Immunol. 2014;71:5:418–26; doi:10.1111/aji.12216.

18. Elovitz MA, Brown AG, Breen K, Anton L, Maubert M, Burd I. Intrauterine inflammation, insufficient to induce parturition, still evokes fetal and neonatal brain injury. Int J Dev Neurosci. 2011;29:6:663–71; doi:10.1016/j.ijdevneu.2011.02.011.

19. Burd I, Bentz AI, Chai J, Gonzalez J, Monnerie H, Le Roux PD, et al. Inflammation-induced preterm birth alters neuronal morphology in the mouse fetal brain. J Neurosci Res. 2010;88:9:1872–81; doi:10.1002/jnr.22368.

20. Elovitz MA, Mrinalini C, Sammel MD. Elucidating the early signal transduction pathways leading to fetal brain injury in preterm birth. Pediatr Res. 2006;59:1:50–5; doi:10.1203/01.pdr.0000191141.21932.b6.

21. Makinson R, Lloyd K, Rayasam A, McKee S, Brown A, Barila G, et al. Intrauterine inflammation induces sex-specific effects on neuroinflammation, white matter, and behavior. Brain Behav Immun. 2017;66:277–88; doi:10.1016/j.bbi.2017.07.016.

22. Rosenkrantz TS, Hussain Z, Fitch RH. Sex Differences in Brain Injury and Repair in Newborn Infants: Clinical Evidence and Biological Mechanisms. Front Pediatr. 2019;7:211; doi:10.3389/fped.2019.00211.

23. Kokras N, Dalla C. Preclinical sex differences in depression and antidepressant response: Implications for clinical research. J Neurosci Res. 2017;95:1–2:731-6; doi:10.1002/jnr.23861.

24. Boivin JR, Piekarski DJ, Wahlberg JK, Wilbrecht L. Age, sex, and gonadal hormones differently influence anxiety-and depression-related behavior during puberty in mice. Psychoneuroendocrinology. 2017;85:78–87; doi:10.1016/j.psyneuen.2017.08.009.

25. Bucci DJ, Chiba AA, Gallagher M. Spatial learning in male and female Long-Evans rats. Behav Neurosci. 2021;135:1:4–7; doi:10.1037/bne0000437.

26. Thagard AS, Slack JL, Estrada SM, Kazanjian AA, Chan S, Burd I, et al. Long-term impact of intrauterine neuroinflammation and treatment with magnesium sulphate and betamethasone: Sex-specific differences in a preterm labor murine model. Sci Rep. 2017;7:1:17883; doi:10.1038/s41598-017-18197-x.

27. Yagi S, Galea LAM. Sex differences in hippocampal cognition and neurogenesis. Neuropsychopharmacology. 2019;44:1:200–13; doi:10.1038/s41386-018-0208-4.

28. Culig L, Surget A, Bourdey M, Khemissi W, Le Guisquet AM, Vogel E, et al. Increasing adult hippocampal neurogenesis in mice after exposure to unpredictable chronic mild stress may counteract some of the effects of stress. Neuropharmacology. 2017;126:179–89; doi:10.1016/j.neuropharm.2017.09.009.

29. Lazarov O, Hollands C. Hippocampal neurogenesis: Learning to remember. Prog Neurobiol. 2016;138-140:1–18; doi:10.1016/j.pneurobio.2015.12.006.

30. Hill AS, Sahay A, Hen R. Increasing Adult Hippocampal Neurogenesis is Sufficient to Reduce Anxiety and Depression-Like Behaviors. Neuropsychopharmacology. 2015;40:10:2368–78; doi:10.1038/npp.2015.85.

31. Revest JM, Dupret D, Koehl M, Funk-Reiter C, Grosjean N, Piazza PV, et al. Adult hippocampal neurogenesis is involved in anxiety-related behaviors. Mol Psychiatry. 2009;14:10:959–67; doi:10.1038/mp.2009.15.

32. Hofer MA, Shair HN, Masmela JR, Brunelli SA. Developmental effects of selective breeding for an infantile trait: the rat pup ultrasonic isolation call. Dev Psychobiol. 2001;39:4:231–46; doi:10.1002/dev.1000.

33. Spijker S. Dissection of Rodent Brain Regions. In: Li K, editor. Neuroproteomics (Neuromethods). Totowa, NJ: Humana Press; 2011. p. 13–26.

34. Warde-Farley D, Donaldson SL, Comes O, Zuberi K, Badrawi R, Chao P, et al. The GeneMANIA prediction server: biological network integration for gene prioritization and predicting gene function. Nucleic Acids Res. 2010;38:Web Server issue:W214–20; doi:10.1093/nar/gkq537.

35. Cristancho AG, Gadra EC, Samba IM, Zhao C, Ouyang M, Magnitsky S, et al. Deficits in seizure threshold and other behaviors in adult mice without gross neuroanatomic injury after late gestation transient prenatal hypoxia. Developmental Neuroscience. 2022; doi:10.1159/000524045.

36. Halekoh U, Højsgaard S, Yan J. TheRPackagegeepackfor Generalized Estimating Equations. Journal of Statistical Software. 2006;15:2:1–11; doi:10.18637/jss.v015.i02.

37. Bates D, Mächler M, Bolker B, Walker S. Fitting Linear Mixed-Effects Models Usinglme4. Journal of Statistical Software. 2015;67:1; doi:10.18637/jss.v067.i01.

38. Caruso A, Ricceri L, Scattoni ML. Ultrasonic vocalizations as a fundamental tool for early and adult behavioral phenotyping of Autism Spectrum Disorder rodent models. Neurosci Biobehav Rev. 2020;116:31–43; doi:10.1016/j.neubiorev.2020.06.011.

39. Jensen FE, Holmes GL, Lombroso CT, Blume HK, Firkusny IR. Age-dependent changes in long-term seizure susceptibility and behavior after hypoxia in rats. Epilepsia. 1992;33:6:971–80; doi:10.1111/j.1528-1157.1992.tb01746.x.

40. Simonet JC, Sunnen CN, Wu J, Golden JA, Marsh ED. Conditional Loss of Arx From the Developing Dorsal Telencephalon Results in Behavioral Phenotypes Resembling Mild Human ARX Mutations. Cereb Cortex. 2015;25:9:2939–50; doi:10.1093/cercor/bhu090.

41. Yardeni T, Cristancho AG, McCoy AJ, Schaefer PM, McManus MJ, Marsh ED, et al. An mtDNA mutant mouse demonstrates that mitochondrial deficiency can result in autism endophenotypes. Proc Natl Acad Sci U S A. 2021;118:6; doi:10.1073/pnas.2021429118.

42. Hester MS, Tulina N, Brown A, Barila G, Elovitz MA. Intrauterine inflammation reduces postnatal neurogenesis in the hippocampal subgranular zone and leads to accumulation of hilar ectopic granule cells. Brain Res. 2018;1685:51–9; doi:10.1016/j.brainres.2018.02.005.

43. Baharnoori M, Bhardwaj SK, Srivastava LK. Neonatal behavioral changes in rats with gestational exposure to lipopolysaccharide: a prenatal infection model for developmental neuropsychiatric disorders. Schizophr Bull. 2012;38:3:444–56; doi:10.1093/schbul/sbq098.

44. Malkova NV, Yu CZ, Hsiao EY, Moore MJ, Patterson PH. Maternal immune activation yields offspring displaying mouse versions of the three core symptoms of autism. Brain Behav Immun. 2012;26:4:607–16; doi:10.1016/j.bbi.2012.01.011.

45. Irie F, Badie-Mahdavi H, Yamaguchi Y. Autism-like socio-communicative deficits and stereotypies in mice lacking heparan sulfate. Proc Natl Acad Sci U S A. 2012;109:13:5052–6; doi:10.1073/pnas.1117881109.

46. Choi GB, Yim YS, Wong H, Kim S, Kim H, Kim SV, et al. The maternal interleukin-17a pathway in mice promotes autism-like phenotypes in offspring. Science. 2016;351:6276:933–9; doi:10.1126/science.aad0314.

47. Fernandez de Cossio L, Guzman A, van der Veldt S, Luheshi GN. Prenatal infection leads to ASD-like behavior and altered synaptic pruning in the mouse offspring. Brain Behav Immun. 2017;63:88–98; doi:10.1016/j.bbi.2016.09.028.

48. Schwartzer JJ, Careaga M, Onore CE, Rushakoff JA, Berman RF, Ashwood P. Maternal immune activation and strain specific interactions in the development of autism-like behaviors in mice. Transl Psychiatry. 2013;3:e240; doi:10.1038/tp.2013.16.

49. Rocha-Gomes A, Teixeira AE, de Oliveira DG, Santiago CMO, da Silva AA, Riul TR, et al. LPS tolerance prevents anxiety-like behavior and amygdala inflammation of high-fat-fed dams’ adolescent offspring. Behav Brain Res. 2021;411:113371; doi:10.1016/j.bbr.2021.113371.

50. Gumusoglu SB, Stevens HE. Maternal Inflammation and Neurodevelopmental Programming: A Review of Preclinical Outcomes and Implications for Translational Psychiatry. Biol Psychiatry. 2019;85:2:107–21; doi:10.1016/j.biopsych.2018.08.008.

51. Hava G, Vered L, Yael M, Mordechai H, Mahoud H. Alterations in behavior in adult offspring mice following maternal inflammation during pregnancy. Dev Psychobiol. 2006;48:2:162–8; doi:10.1002/dev.20116.

52. Smith SE, Li J, Garbett K, Mirnics K, Patterson PH. Maternal immune activation alters fetal brain development through interleukin-6. J Neurosci. 2007;27:40:10695–702; doi:10.1523/JNEUROSCI.2178-07.2007.

53. Wang H, Meng XH, Ning H, Zhao XF, Wang Q, Liu P, et al. Age-and gender-dependent impairments of neurobehaviors in mice whose mothers were exposed to lipopolysaccharide during pregnancy. Toxicol Lett. 2010;192:2:245–51; doi:10.1016/j.toxlet.2009.10.030.

54. Shi L, Fatemi SH, Sidwell RW, Patterson PH. Maternal influenza infection causes marked behavioral and pharmacological changes in the offspring. J Neurosci. 2003;23:1:297–302; https://www.ncbi.nlm.nih.gov/pubmed/12514227.

55. Meyer U. Prenatal poly(i:C) exposure and other developmental immune activation models in rodent systems. Biol Psychiatry. 2014;75:4:307–15; doi:10.1016/j.biopsych.2013.07.011.

56. Meyer U, Nyffeler M, Engler A, Urwyler A, Schedlowski M, Knuesel I, et al. The time of prenatal immune challenge determines the specificity of inflammation-mediated brain and behavioral pathology. J Neurosci. 2006;26:18:4752–62; doi:10.1523/JNEUROSCI.0099-06.2006.

57. Lei Y, Chen CJ, Yan XX, Li Z, Deng XH. Early-life lipopolysaccharide exposure potentiates forebrain expression of NLRP3 inflammasome proteins and anxiety-like behavior in adolescent rats. Brain Res. 2017;1671:43–54; doi:10.1016/j.brainres.2017.06.014.

58. Hsiao EY, McBride SW, Chow J, Mazmanian SK, Patterson PH. Modeling an autism risk factor in mice leads to permanent immune dysregulation. Proc Natl Acad Sci U S A. 2012;109:31:12776–81; doi:10.1073/pnas.1202556109.

59. Depino AM. Early prenatal exposure to LPS results in anxiety-and depression-related behaviors in adulthood. Neuroscience. 2015;299:56–65; doi:10.1016/j.neuroscience.2015.04.065.

60. Dada T, Rosenzweig JM, Al Shammary M, Firdaus W, Al Rebh S, Borbiev T, et al. Mouse model of intrauterine inflammation: sex-specific differences in long-term neurologic and immune sequelae. Brain Behav Immun. 2014;38:142–50; doi:10.1016/j.bbi.2014.01.014.

61. Brown AG, Tulina NM, Barila GO, Hester MS, Elovitz MA. Exposure to intrauterine inflammation alters metabolomic profiles in the amniotic fluid, fetal and neonatal brain in the mouse. PLoS One. 2017;12:10:e0186656; doi:10.1371/journal.pone.0186656.

62. Eichenbaum H, Sauvage M, Fortin N, Komorowski R, Lipton P. Towards a functional organization of episodic memory in the medial temporal lobe. Neurosci Biobehav Rev. 2012;36:7:1597–608; doi:10.1016/j.neubiorev.2011.07.006.

63. Gaesser B, Spreng RN, McLelland VC, Addis DR, Schacter DL. Imagining the future: evidence for a hippocampal contribution to constructive processing. Hippocampus. 2013;23:12:1150–61; doi:10.1002/hipo.22152.

64. Schacter DL, Addis DR, Buckner RL. Remembering the past to imagine the future: the prospective brain. Nat Rev Neurosci. 2007;8:9:657–61; doi:10.1038/nrn2213.

65. Aimone JB, Li Y, Lee SW, Clemenson GD, Deng W, Gage FH. Regulation and function of adult neurogenesis: from genes to cognition. Physiol Rev. 2014;94:4:991–1026; doi:10.1152/physrev.00004.2014.

66. Gu Y, Arruda-Carvalho M, Wang J, Janoschka SR, Josselyn SA, Frankland PW, et al. Optical controlling reveals time-dependent roles for adult-born dentate granule cells. Nat Neurosci. 2012;15:12:1700–6; doi:10.1038/nn.3260.

67. Kohman RA, Rhodes JS. Neurogenesis, inflammation and behavior. Brain Behav Immun. 2013;27:1:22–32; doi:10.1016/j.bbi.2012.09.003.

68. Lucassen PJ, Oomen CA, Naninck EF, Fitzsimons CP, van Dam AM, Czeh B, et al. Regulation of Adult Neurogenesis and Plasticity by (Early) Stress, Glucocorticoids, and Inflammation. Cold Spring Harb Perspect Biol. 2015;7:9:a021303; doi:10.1101/cshperspect.a021303.

69. Kitabatake Y, Sailor KA, Ming GL, Song H. Adult neurogenesis and hippocampal memory function: new cells, more plasticity, new memories? Neurosurg Clin N Am. 2007;18:1:105–13, x; doi:10.1016/j.nec.2006.10.008.

70. Lieberwirth C, Pan Y, Liu Y, Zhang Z, Wang Z. Hippocampal adult neurogenesis: Its regulation and potential role in spatial learning and memory. Brain Res. 2016;1644:127–40; doi:10.1016/j.brainres.2016.05.015.

71. Kelley MH, Wu WW, Lei J, McLane M, Xie H, Hart KD, et al. Functional changes in hippocampal synaptic signaling in offspring survivors of a mouse model of intrauterine inflammation. J Neuroinflammation. 2017;14:1:180; doi:10.1186/s12974-017-0951-1.

72. Ng PC, Wong SP, Chan IH, Lam HS, Lee CH, Lam CW. A prospective longitudinal study to estimate the “adjusted cortisol percentile” in preterm infants. Pediatr Res. 2011;69:6:511–6; doi:10.1203/PDR.0b013e31821764b1.

73. Watterberg KL, Scott SM, Naeye RL. Chorioamnionitis, cortisol, and acute lung disease in very low birth weight infants. Pediatrics. 1997;99:2:E6; doi:10.1542/peds.99.2.e6.

74. Kim JJ, Diamond DM. The stressed hippocampus, synaptic plasticity and lost memories. Nat Rev Neurosci. 2002;3:6:453–62; doi:10.1038/nrn849.

75. Almela M, Hidalgo V, van der Meij L, Pulopulos MM, Villada C, Salvador A. A low cortisol response to acute stress is related to worse basal memory performance in older people. Front Aging Neurosci. 2014;6:157; doi:10.3389/fnagi.2014.00157.

76. Boksa P. Effects of prenatal infection on brain development and behavior: a review of findings from animal models. Brain Behav Immun. 2010;24:6:881–97; doi:10.1016/j.bbi.2010.03.005.

77. Giovanoli S, Notter T, Richetto J, Labouesse MA, Vuillermot S, Riva MA, et al. Late prenatal immune activation causes hippocampal deficits in the absence of persistent inflammation across aging. J Neuroinflammation. 2015;12:221; doi:10.1186/s12974-015-0437-y.

78. Golan HM, Lev V, Hallak M, Sorokin Y, Huleihel M. Specific neurodevelopmental damage in mice offspring following maternal inflammation during pregnancy. Neuropharmacology. 2005;48:6:903–17; doi:10.1016/j.neuropharm.2004.12.023.

79. Khan D, Fernando P, Cicvaric A, Berger A, Pollak A, Monje FJ, et al. Long-term effects of maternal immune activation on depression-like behavior in the mouse. Transl Psychiatry. 2014;4:e363; doi:10.1038/tp.2013.132.

80. Solek CM, Farooqi N, Verly M, Lim TK, Ruthazer ES. Maternal immune activation in neurodevelopmental disorders. Dev Dyn. 2018;247:4:588–619; doi:10.1002/dvdy.24612.

81. Glass R, Norton S, Fox N, Kusnecov AW. Maternal immune activation with staphylococcal enterotoxin A produces unique behavioral changes in C57BL/6 mouse offspring. Brain Behav Immun. 2019;75:12–25; doi:10.1016/j.bbi.2018.05.005.

82. Jin J, Kim SN, Liu X, Zhang H, Zhang C, Seo JS, et al. miR-17-92 Cluster Regulates Adult Hippocampal Neurogenesis, Anxiety, and Depression. Cell Rep. 2016;16:6:1653–63; doi:10.1016/j.celrep.2016.06.101.

83. Goncalves JT, Schafer ST, Gage FH. Adult Neurogenesis in the Hippocampus: From Stem Cells to Behavior. Cell. 2016;167:4:897–914; doi:10.1016/j.cell.2016.10.021.

